# Synchronous Hatching of 2-Day Delayed Intruders in Reproductive Competition in the Burying Beetle

**DOI:** 10.1101/2025.05.16.654448

**Authors:** Takuma Niida, Tomoyosi Nisimura

**Author notes:** Corresponding Author: T. Nisimura, College of Bioresource Sciences, Nihon University, 1866 Kameino, Fujisawa, Japan. T. Niida, Graduate School of Agriculture, Kyoto University, Kitashirakawa-Oiwake-cho, Sakyo-ku, Kyoto, Japan.

## Abstract

Appropriate hatching time is important for successful brood parasitism in some bird and insect groups. Parental burying beetles typically rear their larvae on carcasses buried in nests; however, they occasionally raise brood-parasitic larvae from eggs dumped by other females. Parasitic larvae must hatch synchronously with host larvae because the host parents indiscriminately accept any larvae during that period. However, parasites in burying beetles are assumed to have more severe restrictions on synchronous hatching than those in birds because of less predictability of reproductive sites, the necessity for ovarian development at the site, and difficulty in adjusting by oviposition. Is the hatching of parasitic larvae synchronized with that of host larvae, even if the reproductive behavior of the parasites lags behind that of the host owing to restrictions? In the burying beetle *Nicrophorus quadripunctatus* Kraatz, we observed the hatching of host and parasite offspring under experimental conditions in which female parasites (intruders) encountered carcasses that had been secured by female hosts (residents) two days earlier. In 14 of the 40 replicates, defeated intruders dumped eggs, residents laid 36 eggs, and intruders laid six eggs on average. The hatching of intruder offspring was synchronized with that of the resident offspring despite the two-day lag. Egg dumping is regarded as brood parasitism. Notably, the period from contact with carcasses to hatching was significantly shorter for intruders than for residents. Advances in hatching time may contribute to the success of brood parasitism.

## Introduction

Intra- or interspecific brood parasitism is a reproductive strategy used by several birds and insects. Parasitic females lay eggs in the nests of either conspecific or heterospecific host parents, who provide parental care (Brockmann, 1993; Hamilton & Orians, 1965; Lyon & Eadie, 2008; Payne, 1977; Tallamy, 2005; Yom-Tov, 1980).

Time is critical in influencing the behavioral interactions between hosts and parasites during various brood parasitism events. In birds, the timing of when parasites lay eggs and hatch during the reproductive process of hosts is important for avoiding several antiparasitic behaviors by hosts and ultimately facilitating successful parasitism (Hauber, 2003; Petrie & Møller, 1991). For example, the early dumping of eggs reduces their acceptance by hosts in the barn swallow, *Hirundo rustica* (Møller, 1987), while late dumping decreases the hatching success of parasites in the moorhen, *Gallinula chloropus* (Gibbons, 1986). Furthermore, the late hatching of parasitic chicks decreases their survival in the shiny cowbird *Molothrus bonariensis* (Fiorini et al., 2009). Similarly, brood parasites in some insects, such as Hymenoptera (Field, 1992) and burying beetles (Eggert & Müller, 2000; Eggert & Müller, 2011; Müller & Eggert, 1990; Trumbo, 1994), require timely egg laying and hatching.

Burying beetles, *Nicrophorus* spp. (Coleoptera: Silphidae: Nicrophorinae), provide parental care for reproduction by utilizing the carcasses of small vertebrates (Scott, 1998). The beetles typically take 1 or 2 days to prepare a carcass for their offspring. This process includes burying, removing feathers, rounding, and applying secretions. Carcasses are unpredictably distributed and ephemeral resources (Tallamy & Wood, 1986; Wilson, 1975), with several adults converging on a single carcass, resulting in aggressive intra- and interspecific competition. Due to intrasexual competition, only one larger female or one larger male-female pair buries a carcass into an underground nest (Bartlett & Ashworth, 1988; Otronen, 1988; Scott, 1998; Trumbo, 1994; Wilson & Fudge, 1984). The females lay eggs in the soil surrounding their carcasses, and the adults feed on and guard their hatched larvae.

Intra- and interspecific brood parasitism have been reported in several burying beetle species (Müller et al., 1990; Smith & Belk, 2018; Trumbo, 1994; Trumbo et al., 2001). Intraspecific brood parasitism has been extensively studied in *Nicrophorus vespilloides* Herbst (Eggert & Müller, 2011; Müller et al., 1990, 2007). In competition for carcasses, some losing females (parasites) do not immediately leave the carcasses secured by winning beetles (hosts); the parasites deposit their eggs into the soil near the nest (carcass), relying on the winning beetles to care for their larvae. Hosts can cull larvae that appear on carcasses before their larvae hatch but accept all larvae indiscriminately after their larvae have hatched. However, late-hatching larvae within a brood in burying beetles tend to have lower body mass, growth rates, and survival rates because of sibling competition (Smiseth et al., 2007). Even if late-hatching intruder larvae are accepted by hosts, they may be outcompeted by host larvae. Therefore, parasites must hatch their eggs synchronously with the host larvae in burying beetles (Richardson et al., 2021).

However, the intruders of burying beetles are assumed to have some restrictions on synchronization rather than those of birds. The location of host nests may be more predictable in birds than in burying beetles because birds often reproduce in the same place and season annually (Hamilton & Orians, 1965; Hildén, 1965; Perrins, 1970). Parasitic birds may have opportunities to avoid detection by their hosts and attempt to lay eggs because of the predictability of host nest occurrence. In contrast, the occurrence of breeding sites is highly unpredictable for burying beetles because their reproduction is dependent on rare and ephemeral carcasses (Tallamy & Wood, 1986; Tallamy, 2005; Wilson, 1975). The discovery of carcasses by adults is stochastic, making it challenging for intruders to anticipate the breeding site of hosts in advance.

Additionally, physiological differences exist in the preparation of birds and burying beetles for oviposition. Although female birds generally prepare for egg laying before a specific breeding season (Hau, 2001; Marshall, 1959; Perrins, 1996), ovarian development in female beetles occurs rapidly on site after locating a carcass and is triggered by subsequent behaviors, such as burying and assessing the suitability of carcasses for reproduction (Scott & Traniello, 1987; Trumbo et al., 1995; Wilson & Knollenberg, 1984). When nests are already established and protected by hosts, access to the carcasses is restricted by the host defense behavior, and ovarian development in the intruders may occur later than that in the hosts.

Moreover, the adjustments in the timing of egg dumping are more difficult in burying beetles than those in birds because of differences in the visibility of eggs laid and the process of embryo development. Parasites encounter more difficulties watching the egg laying behavior of hosts in burying beetles that nest and oviposit underground than in birds that typically build terrestrial and visible nests. In birds, embryo development begins synchronously within broods through parental incubation. Therefore, when parasites dump eggs in the period before incubation by hosts, the timing of hatching of parasitic eggs is generally synchronous with that of the host eggs (Slagsvold, 1986; Stoleson & Beissinger, 1995; White & Kinney, 1974). In burying beetles, as well as other insects, embryos develop when eggs are laid because embryo development is generally dependent on environmental temperatures (Takata et al., 2013; Tallamy, 2005). Therefore, precise synchronization of egg laying between hosts and parasites may be required for synchronous embryo development.

Considering these restrictions, intruders are expected to hatch their eggs less synchronously with their resident eggs. Despite these restrictions, intra- and inter-brood parasitism has been reported in burying beetles (Müller et al., 2007; Müller et al., 1990; Smith & Belk, 2018; Trumbo et al., 2001; Trumbo, 1994). In specific processes in brood parasitism, female beetles locate nests previously secured by other females in the field (Robertson, 1993; Scott, 1990; Trumbo, 1990a). Under laboratory conditions involving *N. pustulatus,* females deposited a few eggs around the carcasses secured 4 days before by a host female (Trumbo & Valletta, 2007). However, whether synchronous hatching between the host and parasitic larvae occurs when females contact the carcass after a time lag remains unclear. Clarifying synchronous hatching between hosts and parasites is important for understanding the mechanisms of brood parasites in burying beetles.

In this study, we investigated egg dumping by intruders (parasites) and egg laying by residents (hosts) in the burying beetle *N. quadripunctatus* Kraatz. The experiments were conducted under conditions where parasitic females invaded the nests that had been secured by host females 2 days earlier. We also examined whether their larvae synchronously hatched.

## METHODS

### Insects and preparation for behavioral experiments

Adults of the burying beetle species *N. quadripunctatus* were collected using hanging traps (15.5 × 15 × 17 cm) in a deciduous forest on the Nihon University College of Bioresource Sciences (NUBS) campus (35° 22’ N, 139° 27’ E) between April and June, September and October 2017, and March, April, and September 2018. The traps were baited with a piece of chicken meat (approximately 50 g) and installed approximately 1 m above the ground on tree trunks. The traps were checked once every 3 days, and 43 males and 46 females were collected. Field-collected adults were then maintained as male-female pairs in plastic cups (9 cm diameter × 4.5 cm height), which were filled with approximately 40 g of soil collected from a NUBS field. The pairs were maintained under a photoperiod of 12 h light and 12 h darkness (LD 12:12 h) and a temperature of 20 ± 1 °C using an incubator (MIR-154-PJ, Panasonic Healthcare Co., Ltd., Tokyo, Japan). These are the non-diapause conditions for *N. quadripunctatus* (Nisimura et al., 2002). All the adults were fed a small piece of chicken liver *ad libitum*.

To obtain laboratory-reared adults (F_1_ generation) for behavioral experiments, pairs were reproduced with approximately 25 g of chicken meat in plastic containers (11.5 cm diameter × 10 cm height) filled with soil to a depth of approximately 4 cm. Larvae that ceased feeding and began wandering were kept for each brood in soil-filled plastic containers (15.5 cm diameter × 9 cm height) to facilitate pupation.

Newly emerged F_1_ adults were kept under the same conditions as field-collected adults; however, some were kept under a photoperiod of LD 14:10, which was also used as a non-diapause condition (Takata et al., 2015). The day that each adult emerged from the soil was recorded as the day of emergence. The width of the pronotum of each adult was measured using digital calipers (DT-100; Niigata Seiki Co., Ltd., Niigata, Japan). Each adult pair was fed 10 fly larvae (*Calliphora nigribarbis* Snellen van Vollenhoven, *Aldrichina graham* Aldrich, *Lucilia Caesar* Linnaeus, *L. illustris* Meigen, or *Muscina stabulans* Fallen) two times for the first week after emergence. Flies were collected from a NUBS field and reared on chicken livers.

For the behavioral experiments, the pairs were fed chicken liver (0.5 g) with or without Sudan red (100–200 mg Sudan red per 20 g liver) (Tokyo Chemical Industry Co., Ltd., Tokyo, Japan) twice a week. Sudan red is a fat-soluble dye that causes adult females to produce pink eggs (Eggert et al., 2008); therefore, it was possible to distinguish which females lay eggs in the experiments.

### Behavioral experiments

To investigate the timing of offspring hatching from intruding and resident females, we observed the reproductive behavior of two adult females that were kept as male-female pairs in a laboratory. The females contacted the resources with a time lag in a plastic container (11.5 cm diameter × 10 cm height). This container was attached to a transparent plastic cup (9 cm diameter × 4.5 cm height), which served as a refuge space for females that were not successful in competition (Fig. 1). The container was filled with soil to a depth of 4 cm, and approximately 25.0 g of chicken meat was placed on the soil surface. Only females reared as male-female pairs were used to exclude the effects of males on the experimental results. We assumed that most females copulated until the experiments because copulation was often observed inside the rearing cups.

**Fig. 1.**
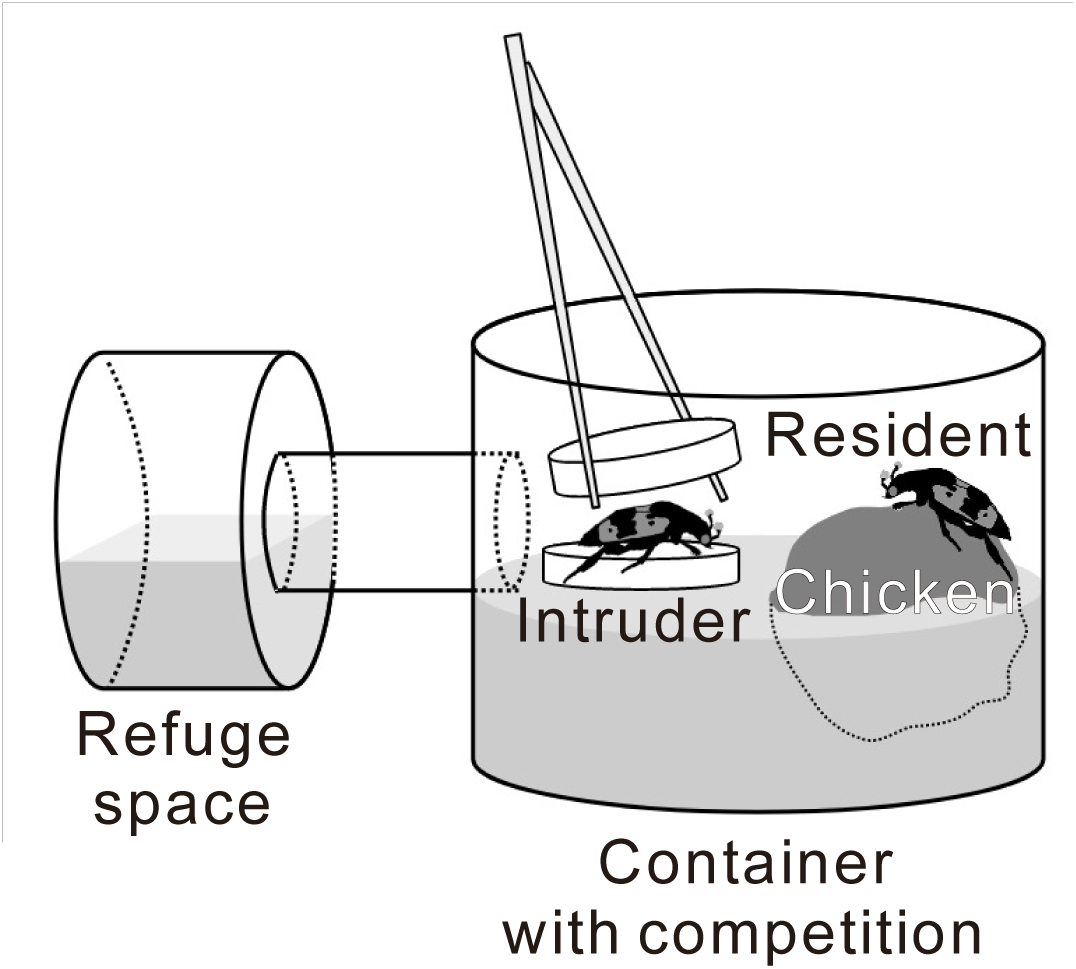
Overview of behavioral experiments under competitive conditions in *Nicrophorus quadripunctatus*. Two days after the residents began reproducing chicken meat in a plastic container, an intruder was introduced into the container. When introduced, the intruder was caged in a plastic Petri dish and placed on the soil in the container. The Petri dish was carefully opened using tweezers to allow the intruder to compete with the residents. The container was attached to a transparent plastic cup that was used as a refuge for the females. See Materials and Methods for further details.

On the first day, one female fed with a non-dyed liver was introduced into a container as a resident female (resident) (Fig. 1). Two days after beginning the experiment, another female fed with a dyed liver was introduced into the container as an intruding female (intruder). To ensure that intruders were disadvantaged in the competition, females who were smaller than the residents were selected as intruders (Suzuki, 2011). To prevent a resident from being disturbed due to panic response of the intruder at induction, the intruder was caged inside a plastic Petri dish (3.5 cm diameter × 1 cm height) and then kept on the soil of the containers for approximately 20 min for acclimatization. The Petri dish was then carefully opened to allow the intruders to compete with the residents (Fig. 1).

The containers and refuges were checked daily. The behavioral experiment ceased when any of the following outcomes occurred:

The resident won: the intruder appeared at the refuge or died.
The intruder won: the resident appeared at the refuge or died.
A winner was not identified: hatched larvae appeared on the chicken meat, and no females were found at the refuge.

Females removed from the containers were identified as either residents or intruders by measuring the width of their pronota.

Behavioral experiments without intruders, namely under non-competitive conditions, were also conducted using a single female fed with either a dyed or non-dyed liver.

### Eggs and larval hatching

After the behavioral experiments, the soil in the container was carefully transferred onto a tray (44 × 32.5 × 7 cm), brushed, and the eggs were subsequently removed. The number of normal-colored eggs (resident offspring) and pink eggs (intruder offspring) were counted. The number of hatched larvae that were observed before the cessation of behavioral experiments was also counted as resident offspring because residents were induced in the experiments 2 days earlier than the intruders.

The eggs were photographed from the top with a ruler using a digital camera (Coolpix aw130; Nikon Corp., Tokyo, Japan) to estimate their volume. The lengths of the major and minor axes of eggs were measured using a ruler. Egg volume (V) was then calculated using the following formula, assuming an egg to be an ellipsoid shape: V (mm³) = 4/3 × π × α/2 × (β/2)^2^, where π is the circumference ratio, and α and β are the lengths of the major and minor axes of the eggs, respectively. The egg volume of each clutch was calculated by averaging the volume of eggs.

The eggs were subsequently kept in an incubator on wet cotton in a plastic case (6.7 × 3.8 × 1.5 cm). The hatched larvae were examined daily until all eggs hatched or died. The hatching day of each egg was recorded from the first day on which a resident or intruder contacted a resource (the hatching day was calculated based on the beginning of the behavioral experiments for residents and two days after the beginning of the experiments for intruders). The hatching day of each resident or intruder clutch was calculated by averaging the hatching days of the eggs.

### Competition duration

In behavioral experiments with competition, the period of coexistence of intruders with residents was estimated as the competitive duration. To examine the duration of time intruders needed to stay near the carcass to allow egg dumping, the competitive duration was compared between cases in which intruders did and did not oviposit.

### Statistical analysis

All statistical analyses were performed using the R software (R Core Team, 2018). Generalized linear mixed models (GLMMs) or linear mixed models (LMMs) were used to analyze differences in hatching day, number of eggs, hatching ability, egg volume, competitive duration, and intruder oviposition (Table A1). Individual identity (individual ID) was used as a random effect for hatching day, hatching ability, and egg volume, and experimental identity (experiment ID) was used for the number of eggs, competitive duration, and intruder oviposition.

In the GLMMs for hatching day, number of eggs, and competitive duration, a Poisson distribution (log-link function) was used as the probability distribution because the response variables were positive discrete numbers. A binomial distribution (logit link function) was used in the GLMMs for hatching ability and intruder oviposition, which were two-valued variables. A Gaussian distribution (identity link function) was used in the LMM for egg volume, which had a positive continuous number.

Age (days from adult emergence), size (width of the pronotum at emergence), arrival order (resident = 0; intruder = 1), competitive condition (non-competitive = 0; competitive condition = 1), and dyed food (non-dyed food = 0; dyed food = 1) were included as explanatory variables in the models for hatching day, number of eggs, hatching ability, and egg volume (Table S1). The age of the residents and intruders, their size, and oviposition by the intruder (not oviposited = 0; oviposited = 1) were included as explanatory variables in the analysis of competitive duration. The ages of the residents and intruders and their sizes were also included as explanatory variables in the model of oviposition by intruders.

GLMMs were constructed using the glmmML package (Broström, 2020), whereas LMMs were constructed using the lme4 (Bates et al., 2015) and lmerTest packages (Kuznetsova et al., 2017).

Moreover, Kruskal–Wallis tests were performed to examine whether the age and size of females differed among the four conditions, that is, residents and intruders under competitive conditions and residents fed non-dyed and dyed food under non-competitive conditions.

## RESULTS

### Reproductive behavior

Residents buried chicken meat for reproduction in 49 of 66 replicates under competitive conditions (Table S2). Intruders were introduced into the containers of 49 replicates: intruders took refuge in 40 replicates (A in the outcome category), residents took refuge in eight replicates (B), and no females took refuge in one replicate (C). Of the 40 replicates in which intruders took refuge (A), normal eggs alone were observed in 19 replicates, pink eggs alone in 1 replicate, normal-colored and pink eggs in 13 replicates, and no eggs in 7 replicates. Therefore, 14 intruders laid their eggs, even though they lost in competition.

Of the 38 replicates under non-competitive conditions, 10 females fed non-dyed food and 10 females fed dyed food buried chicken meats and subsequently oviposited

(Table S2).

In subsequent analyses, we focused on 40 replicates in which intruders lost under competitive conditions (A) and 20 replicates in which reproductive behavior was observed under non-competitive conditions.

### Age and size of females used in behavioral experiments

In the 40 replicates under competitive conditions, the mean age at the beginning of experiments was 36.4 ± 12.9 (± SD) days after emergence for residents (n = 40) and 39.2 ± 13.3 days for intruders (n = 40). The mean pronotum width was 5.80 ± 0.15 mm for residents (n = 40) and 5.29 ± 0.28 mm for intruders (n = 40).

In the 20 replicates under non-competitive conditions, the mean age at the beginning of experiments was 31.8 ± 7.77 days after emergence in females fed non-dyed food (n = 10) and 36.7 ± 7.44 days in females fed dyed food (n = 10). The mean pronotum width was 5.85 ± 0.11 mm in females fed non-dyed food (n = 10) and 5.49 ± 0.57 mm in females fed dyed food (n = 10).

Among females under these conditions (residents and intruders under competitive conditions and residents fed non-dyed and dyed food under non-competitive conditions), no significant differences were observed in the mean age (Kruskal–Wallis test: chi-square = 2.8599, df = 3, P > 0.05), whereas significant differences were observed in the mean pronotum width due to the choice of smaller individuals as intruders (Kruskal–Wallis test: chi-squared = 56.381, df = 3, P < 0.05).

### Hatching day

Of the 14 replicates in which intruders dumped eggs, two were removed from residents: one resident did not lay eggs in one replicate, and the origin of the larvae was not identified in the other replicate because we confused the hatched larvae derived from a resident or intruder. The mean hatching day for 12 residents was 6.08 ± 1.69 days (n = 12) after the beginning of experiments (Fig. 2). Of the 14 replicates, four were removed from intruders: the origin of the larvae was not identified in one replicate, and no larvae hatched in three replicates. The mean hatching day for 10 intruders was 4.57 ± 0.61 days (n = 10) after their introduction. Of the 10 replicates under non-competitive conditions, the mean hatching day for 10 residents fed non-dyed food was 5.30 ± 1.03 days (n = 10) after the beginning of experiments. Of the 10 replicates, three were removed from residents fed dyed food because there were no hatched larvae. The mean hatching day for seven residents fed dyed food was 5.72 ± 1.06 days (n = 7) after the beginning of experiments.

**Fig. 2.**
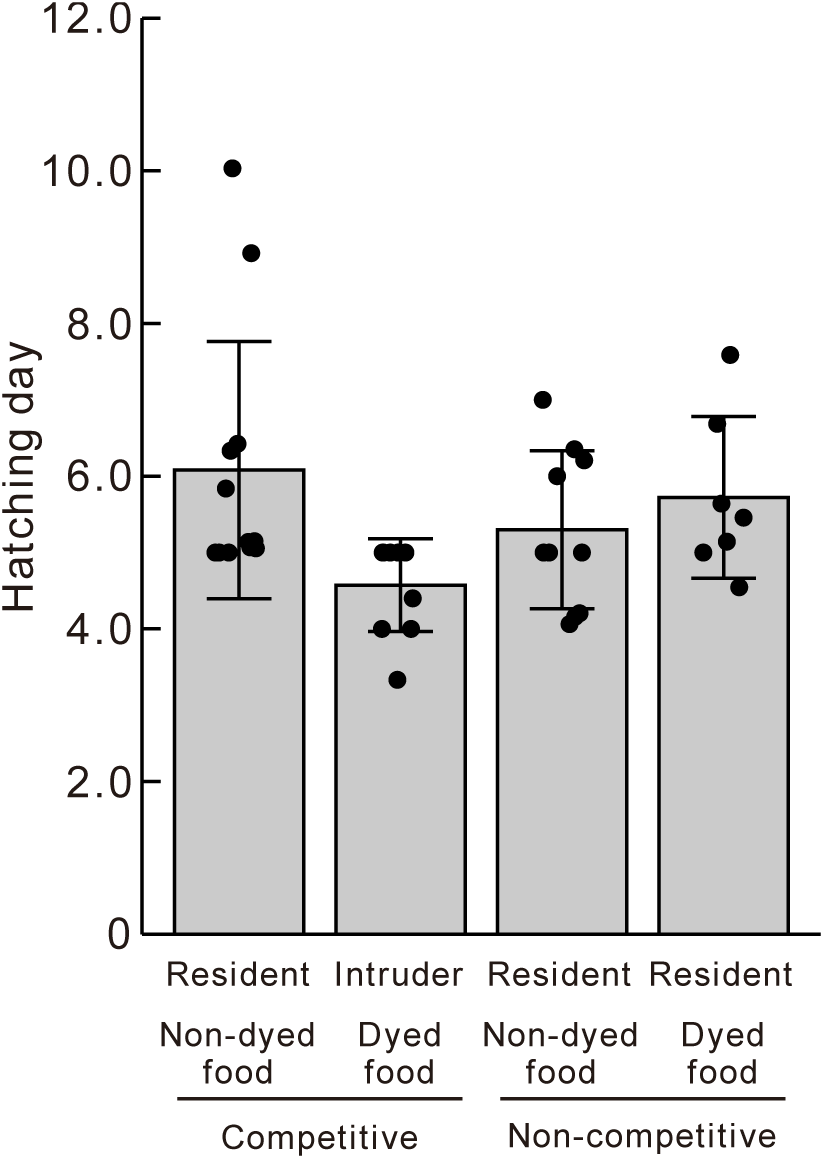
Hatching day of *N. quadripunctatus*. Black points show the average hatching day of each individual, which are residents and intruders under competitive and non-competitive conditions (n = 12, 10, 10, 7). Gray and error bars represent the mean and SD of the average hatching day.

The GLMM analysis showed whether a female was a resident or intruder; the arrival order (resident = 0; intruder = 1) negatively correlated with the hatching day (Table S3; GLMM, P <0.05). Competitive conditions and dyed food did not significantly affect hatching days (Table S3; GLMM, P >0.05). Therefore, the period from contact with resources to hatching was significantly shorter for the intruders than for the residents.

### Number of eggs

Under competitive conditions, the mean number of eggs was 35.5 ± 15.8 (n = 14) for 14 residents and 6.07 ± 4.12 (n = 14) for 14 intruders (Fig. 3). Under non- competitive conditions, the mean number of eggs was 35.3 ± 12.7 (n = 10) for 10 residents fed non-dyed food and 35.1 ± 12.9 (n = 10) for 10 residents fed dyed food.

**Fig. 3.**
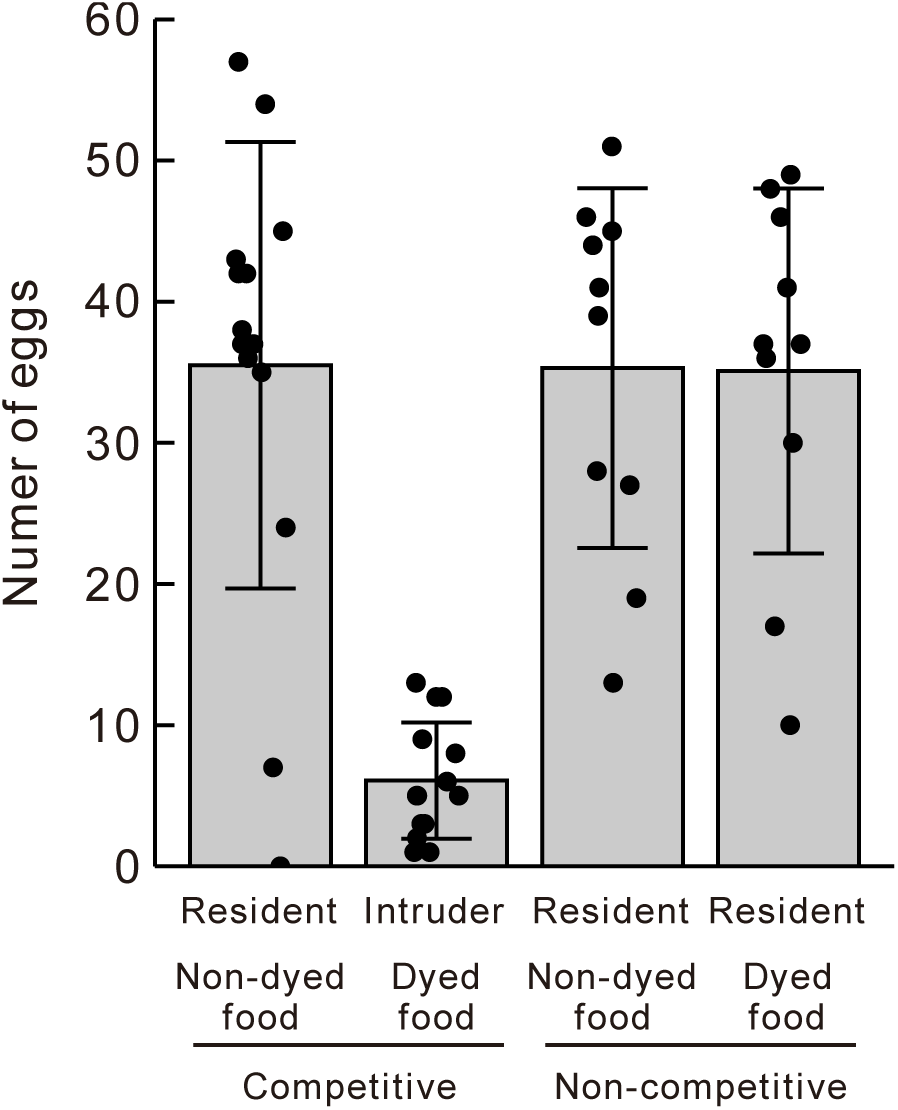
The number of eggs of *N. quadripunctatus.* Black points show the number of eggs of each individual, which are residents and intruders under competitive and non-competitive conditions (n = 14, 14, 10, 10). Gray and error bars represent the mean and SD of the number of eggs.

GLMM analysis showed that only the arrival order significantly affected the number of eggs (Table S4; GLMM, P <0.05). Therefore, the number of eggs was lower for the intruders than those for the residents.

### Hatching rate

Under competitive conditions, the mean hatching rate of 13 residents was 84.2% (n = 13) (Fig. 4). One replicate in which the resident did not oviposit was excluded from the analysis, whereas one replicate in which the origin of the larvae was not identified was included in the hatching rate analysis because all the eggs hatched. The mean hatching rate of the 14 intruders was 56.0% (n = 14). Under non-competitive conditions, the mean hatching rates of 10 residents fed non-dyed food and 10 residents fed dyed food were 68.4 (n = 10) and 48.4% (n = 10), respectively.

**Fig. 4.**
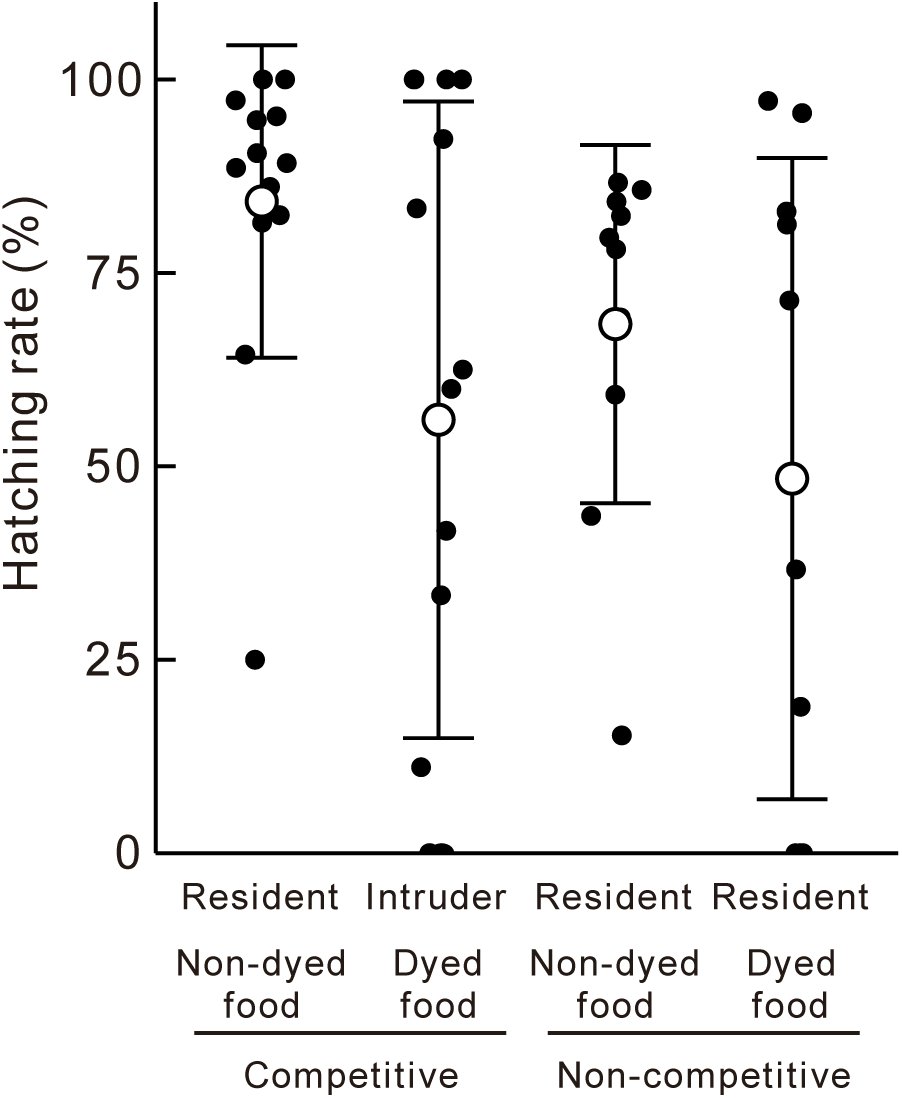
Hatching rate of *N. quadripunctatus*. Black points show the hatching rate of each individual, which are residents and intruders under competitive and non-competitive conditions (n = 13, 14, 10, 10). Large white points and error bars represent the mean and SD of the hatching rate.

GLMM analysis showed that no explanatory variables had a significant effect on hatching ability (Table S5; GLMM, P >0.05).

### Egg volume

Under competitive conditions, the mean egg volume for the 13 residents was 2.11 ± 0.28 mm³ (n = 13) (Fig. 5), although one replicate was removed because no eggs were observed. Of the 497 eggs obtained from the residents, 76 (0–34 eggs contained in each replicate) were also excluded from the analysis, and accurate measurements were challenging because the soil adhered to the eggs. The mean egg volume for the 14 intruders was 1.75 ± 0.48 mm³ (n = 14). Of the 85 eggs obtained from the intruders, four (0–2 eggs contained in each replicate) were excluded from the analysis because the soil adhered to the eggs.

**Fig. 5.**
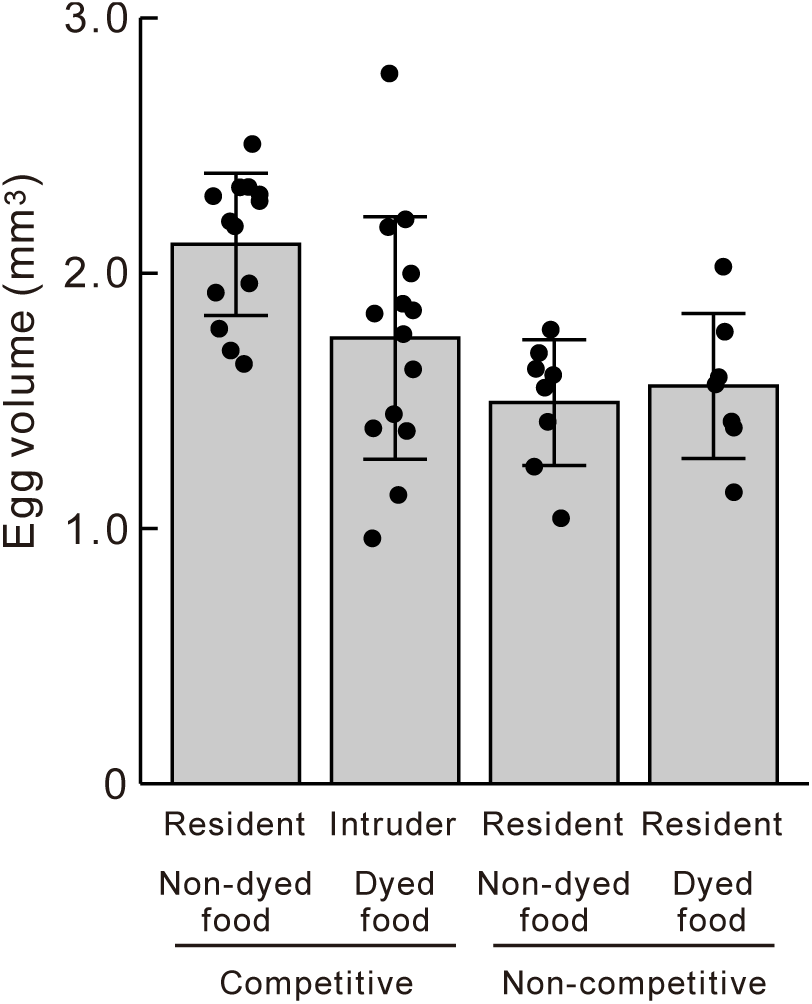
Egg volume of *N. quadripunctatus*. Black points show the average egg volume of each individual, which are residents and intruders under competitive and non-competitive conditions (n = 13, 14, 8, 7). Gray and error bars represent the mean and SD of the average egg volume.

Under non-competitive conditions, the mean egg volume for the eight residents fed non-dyed food was 1.49 ± 0.25 mm³ (n = 8), although two replicates were removed because eggs failed to be photographed. Of the 307 eggs obtained from eight residents fed non-dyed food, 40 (0–39 eggs contained in replicate) were excluded from the analysis because the soil adhered to the eggs. The mean egg volume in seven residents fed dyed food was 1.56 ± 0.28 mm³ (n = 7), although three replicates were excluded from the analysis owing to failure in photographing the eggs. Of the 274 eggs obtained from seven residents fed the dyed food, 14 (0–9 eggs contained in each replicate) were also excluded from the analysis because the soil adhered to the eggs.

The LMM analysis also showed that the arrival order and competitive conditions were both correlated with egg volume (Table S6; LMM, P <0.05). Therefore, intruders oviposited smaller eggs than residents. Furthermore, females oviposited larger eggs under competitive conditions than under non-competitive conditions.

### Competitive duration and oviposition by intruders

In the 40 replicates in which intruders lost under competitive conditions (category A, Table S2), 14 intruders that dumped eggs coexisted alongside residents for 2.57 ± 1.28 days (n = 14) (Fig. 6). Furthermore, 26 intruders that did not oviposit coexisted with residents for 1.69 ± 0.79 days (n = 26). The GLMM analysis showed that no explanatory variables significantly affected competitive duration (Table S7; GLMM, P > 0.05).

**Fig. 6.**
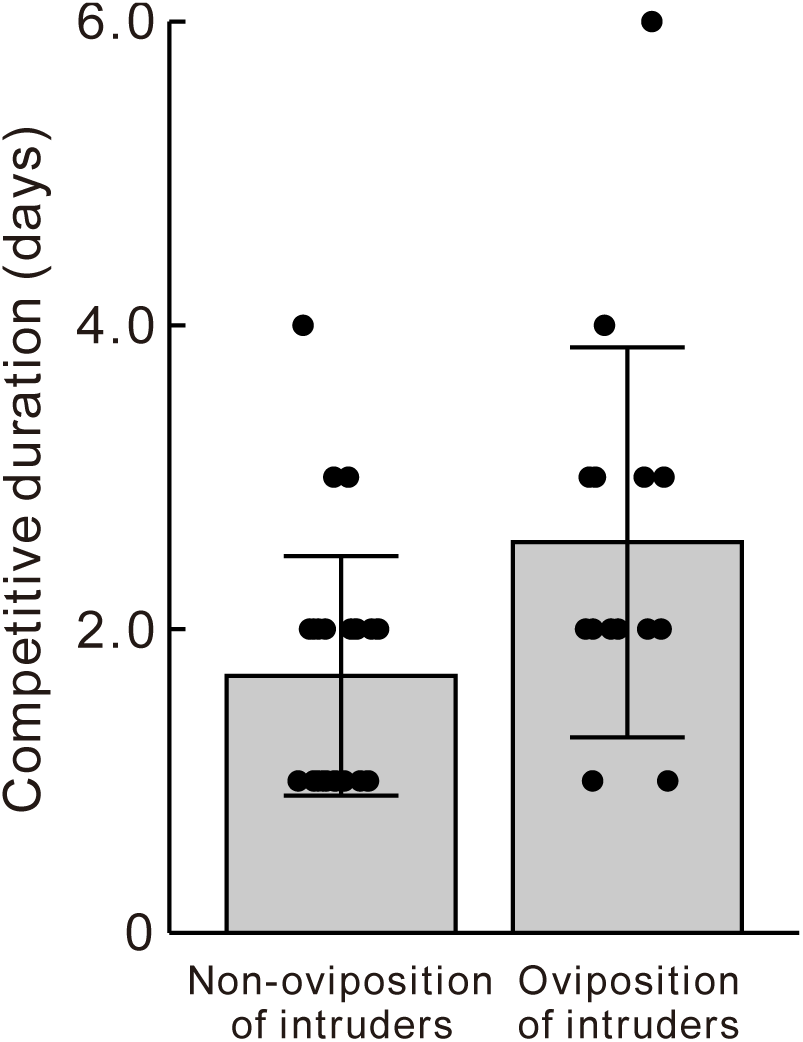
Competitive duration of *N. quadripunctatus*. Black points show the competitive duration of each experiment, which shows non-intruder oviposition and intruder oviposition under competitive conditions (n = 26, 14). Gray and error bars represent the mean and SD of the competitive duration.

The mean age of residents at the beginning of experiments was 36.6 ± 11.3 days (n = 14) when intruders dumped eggs and 36.2 ± 13.9 days (n = 26) when they did not.

The mean ages of the 14 intruders that dumped eggs and the 26 intruders that did not were 40.7 ± 11.8 (n = 14) and 38.4 ± 14.1 days (n = 26), respectively.

The mean pronotum width of residents was 5.75 ± 0.14 mm (n = 14) when intruders dumped eggs and 5.83 ± 0.16 mm (n = 26) when they did not. The mean pronotum widths of the 14 intruders that dumped eggs and the 26 intruders that did not were 5.17 ± 0.35 (n = 14) and 5.35 ± 0.22 mm (n = 26), respectively.

No significant differences were observed in age or pronotum width between intruders that oviposited and those that did not (Table S8; GLMM, P >0.05).

## DISCUSSION

### Egg laying by intruders

In the present study, 14 intruders laid eggs in the soil near chicken meat that residents had utilized. The intruders dumped fewer and smaller eggs than the residents (Figs. 3 and 4); however, no significant difference was observed in hatching ability between residents and intruders (Fig. 5), and the intruder eggs subsequently hatched. If behavioral experiments had continued, the larvae of intruders might have aggregated with those of residents within nests and been accepted by residents because of indiscriminate parental care by females after the hatching of their larvae (Eggert & Müller, 2011). Therefore, we considered egg dumping to function as intraspecific brood parasitism in *N. quadripunctatus*. Parental females of this species occasionally care for genetically unrelated larvae within their broods under natural conditions (Niida et al., 2024), which provides evidence of intraspecific brood parasitism. However, further studies are needed to examine whether hosts sufficiently care for parasitic larvae until they complete larval development and are dispersed from the nests for pupation.

Other reproductive behaviors, other than brood parasitism, should be considered when examining egg-laying by non-resident females in burying beetles. In some *Nicrophorus* species, multiple females engage in cooperative breeding, a reproductive system in which more than one pair of individuals shows parental care towards the young of a single nest or brood (Trumbo & Wilson, 1993). This phenomenon is also referred to as joint (Eggert & Müller, 1992), communal (Trumbo, 1992), communally (Scott & Williams, 1993), or co-breeding (Eggert & Müller, 2000). However, cooperative breeding has not been reported for *N. quadripunctatus*. In the present study, most of the losing females in competition were refuged prior to the appearance of the larvae, and no coexisting of two females on chicken meat were observed. Therefore, it is unlikely that egg laying by intruders was caused by cooperative breeding.

Regarding the traits that caused egg dumping, the age and body size of the residents and intruders did not significantly affect egg dumping (Table S8). Therefore, other physiological traits associated with the losing females may contribute to egg dumping by *N. quadripunctatus*.

Competition duration from their introduction to a refuge was longer for intruders that dumped eggs than for those that did not dump, that is, 2.57 and 1.69 days, respectively (Fig. 6). Intruders require approximately 2–3 days to finish dumping eggs around nests established by residents. Females generally begin to oviposit 12–48 h after carcass discovery in *N. tomentosus* (Scott, 1997, 1998), 10–20 h in *N. vespilloides* (Eggert & Müller, 2000; Richardson & Smiseth, 2020), and approximately 20 h in *N. quadripunctatus* (Takata et al., 2015). Considering the reproductive ecology of *N. quadripunctatus*, a competition duration of 2 or 3 days appears to be sufficient for intruders to dump eggs in *N. quadripunctatus*.

### Synchronization of the hatching day between residents and intruders

The hatching day based on the day when residents and intruders first made contact with chicken meat was earlier for intruders than that for residents (Fig. 2). However, when 2 days were added to the hatching day of the intruders based on the day on which residents contacted the carcasses, no significant difference was observed between the hatching days of residents and intruders (Table S9). This result indicates that the 2-day difference between residents and intruders at the time of contact with reproductive resources was nullified by earlier hatching of intruder larvae. Consequently, larval hatching may have been synchronized between residents and intruders under competitive conditions.

In burying beetles, synchronized hatching between resident and intruder females is generally regarded as a distinct adaptive trait of brood parasitism. Host females (residents) indiscriminately accept *Nicrophorus* larvae after their larvae hatch but kill any larvae that arrive before that time (Müller & Eggert, 1990; Trumbo, 1994). Furthermore, even if late-hatched parasitic larvae are accepted by hosts and provided with parental care, they are expected to be outcompeted by sibling competition. Later-hatching larvae typically have a lower body mass, growth rate, and survival rate than those of earlier hatching larvae and are culled by parents during parental care to match the brood size to resource availability in *N. tomentosus*, *N. vespilloides*, and *N. quadripunctatus* (Bartlett & Ashworth, 1988; Bartlett, 1987; Smiseth et al., 2007; Takata et al., 2013; Trumbo, 1990b). In terms of these restrictions, the ability to cancel differences in hatching times may be a beneficial trait for intruders who find nests already established by the residents of burying beetles.

Variability in hatching time has been reported in previous studies on intraspecific brood parasitism and cooperative breeding of burying beetles. In *N. vespilloides*, the dominant females occasionally avoid synchronous hatching by delaying oviposition, which indicates antiparasitism. The larvae of dominants (hosts) that monopolize carcasses hatch approximately half a day after the larvae of subordinates (parasites), and the larvae of subordinates consequently hatch during the infanticide phase before the hatching of dominant larvae and are eliminated by dominants (Eggert & Müller, 2011; Richardson et al., 2021). In cooperative breeding, even females that coexist on a carcass and jointly care for larvae tend to delay their oviposition and hatching time to reduce the risk of their larvae being killed by cobreeders and eliminate cobreeder larvae (Eggert & Müller, 2000). In contrast to previous findings of delays in the hatching time of host larvae to avoid parasitism, advancing the hatching time of parasitic larvae appeared to promote parasitism. In the intraspecific brood parasitism of insects and birds, advances in the hatching time of parasite offspring for synchronous hatching with host offspring were not widely reported, except for a few studies, such as those on the cliff swallow *Petrochelidon pyrrhonota* (Brown, 1984; Petrie & Møller, 1991).

Compared to birds, intraspecific brood parasites in burying beetles should adapt to the severe restrictions associated with synchronous hatching with hosts: 1) less predictability of reproductive sites, 2) necessity of ovarian development at the site, and 3) difficulty in adjusting by oviposition. Traits by parasitic females to adjust hatching times, even within a short period after locating the nests of other females, may be effective and significant for success in brood parasitism under these restrictions on burying beetles.

The physiological mechanisms underlying the advances in the hatching time of intruder larvae may be based on the smaller eggs of intruders compared to those of residents (Fig. 5). The process from contact with reproductive resources to larval hatching in intruders is divided into two physiological stages: ovarian and embryonic development. One potential mechanism is to advance the timing of oviposition by increasing the rate or early completion of ovarian development, which shortens the ovarian developmental period and may contribute to the small eggs of intruders (Fig. 5). However, egg size is affected by various factors, such as body size in *N. quadripunctatus* (Takata et al., 2015) and the circumstances of co-breeding in *N. vespilloides* (Richardson & Smiseth, 2020).

Therefore, we cannot conclude that this is likely. Furthermore, the period of embryonic development of intruder eggs may be shorter than that of resident eggs. However, in *N. quadripunctatus*, egg volume is not associated with the period of embryonic development (Takata et al., 2015) even though it is positively correlated with the period of embryonic development across several insects (Gillooly & Dodson, 2000; Maino et al., 2017). The eggs of single females under non-competitive conditions were smaller than those of resident females (Fig. 5) but hatched synchronously with those of resident females (Fig. 2). Therefore, there is no evidence of a relationship between small egg volume and advanced hatching times. However, untested physiological mechanisms may be related to adjustments in intruder larval hatching.

Moreover, the behavioral mechanism underlying the advances in hatching time may be assumed as follows. First, the intruder may not take time to prepare a carcass for nesting, whereas the resident must. Second, intruders may require less time to finish laying eggs than residents because intruders oviposit less than residents (Fig. 3). In *N. quadripunctatus*, females take approximately 60 h to lay approximately 20 eggs when provided with 4 g of mouse carcasses (Takata et al., 2015). In this study, the differences in the oviposition period may have caused advances in hatching time because the intruders stopped laying earlier than the residents. In future, to clarify the physiological and behavioral mechanism needs to observe the period of ovary development, oviposition and embryo development, as in Trumbo and Valletta (2007).

Nevertheless, synchronous hatching between intruder and resident may be a coincidence because condition in induction of intruders was limited to 2 days lag in this study. In other words, intruder larval hatch might be advanced a constant period regardless of days on which residents established the nest. In future, therefore, systematic experiments about time lag such as 3 or 4 days should be conducted in order to clarify whether the advancing hatching time of intruders has plasticity depending on circumstances.

In this study, we demonstrated that *N. quadripunctatus* engages in reproductive competition in which losing intruders laid eggs in the nests of residents; however, previous studies have not investigated egg dumping by intruders in burying beetles (Müller et al. 1990; Eggert & Müller, 2000, 2011; Richardson et al., 2021). The hatching time of eggs from intruders was earlier and appeared to synchronize with that of eggs from residents. The advancement in hatching time may be one of the adaptive traits of intruders (parasites) that facilitates successful brood parasitism under competitive association, in which parasites intrude into nests already occupied by residents (hosts) in burying beetles.

## Statements and Declarations

### Ethical note

For the use of animals in research, this study followed the Animal Experiment Manuals by Nihon University Animal Care and Use Committee. *N. quadripunctatus* and some flies are not on the list of threatened species and require no license for research in Japan.

## Acknowledgments

We thank Hidetoshi Iwano, Yoshinori Hatakeyama, and Yuuichi Yamamoto at the Laboratory of Applied Entomology at NUBS; Hiroshi Abé and Hirohiko Takeuchi at the Biological Laboratory at NUBS for providing us with the opportunity to conduct the present study; and Kengo Noma, Nagisa Tosano, Masafumi Hasegawa, Kohei Seto, and Natsumi Katsube for their discussions. We thank Seizi Suzuki for his advice regarding this study.

## Author Contributions

Takuma Niida wrote the main manuscript text, collected data, analyzed data and prepared figures and tables. Tomoyosi Nisimura helped with statistical analysis and figure preparation and edited and revised the manuscript.

## Data Availability

Data are available upon request from authors.

## Supplementary Information (SI)

**Table S1.**
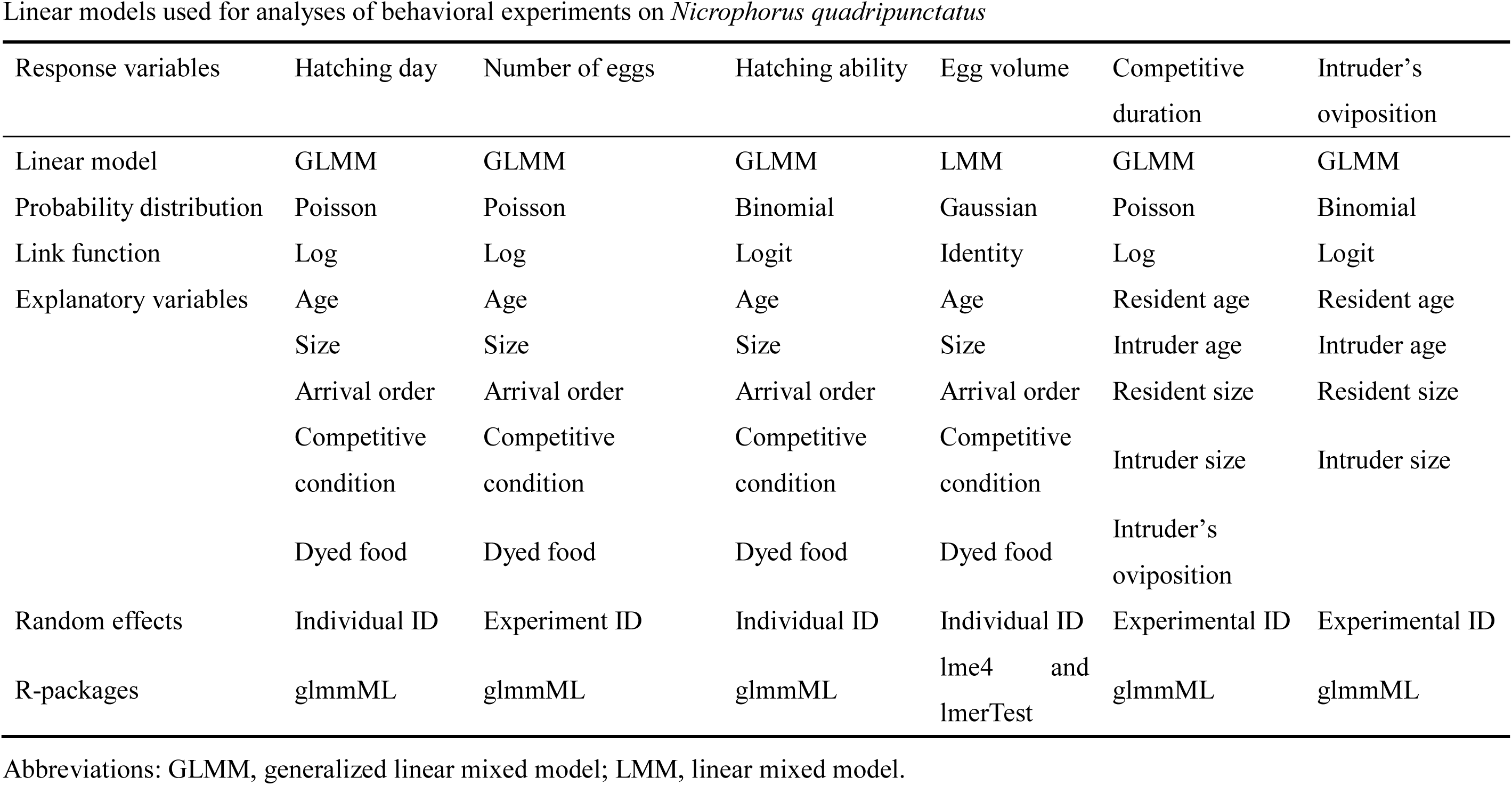
Linear models used for analyses of behavioral experiments on *Nicrophorus quadripunctatus*.

| Response variables | Hatching day | Number of eggs | Hatching ability | Egg volume | Competitive duration | Intruder's oviposition |
| --- | --- | --- | --- | --- | --- | --- |
| Linear model | GLMM | GLMM | GLMM | LMM | GLMM | GLMM |
| Probability distribution | Poisson | Poisson | Binomial | Gaussian | Poisson | Binomial |
| Link function | Log | Log | Logit | Identity | Log | Logit |
| Explanatory variables | Age | Age | Age | Age | Resident age | Resident age |
|  | Size | Size | Size | Size | Intruder age | Intruder age |
|  | Arrival order | Arrival order | Arrival order | Arrival order | Resident size | Resident size |
|  | Competitive condition | Competitive condition | Competitive condition | Competitive condition | Intruder size | Intruder size |
|  | Dyed food | Dyed food | Dyed food | Dyed food | Intruder's oviposition |  |
| Random effects | Individual ID | Experiment ID | Individual ID | Individual ID | Experimental ID | Experimental ID |
| R-packages | glmmML | glmmML | glmmML | lme4 and lmerTest | glmmML | glmmML |

Abbreviations: GLMM, generalized linear mixed model; LMM, linear mixed model.

**Table S2.** Frequency of reproductive behavior and oviposition by the burying beetle *N. quadripunctatus* reared under 12 h light and 12 h darkness (LD 12:12 h) conditions and 14:10 h at 20 °C. a) The categories of outcomes A, B, and C are described in detail in the Materials and Methods.

|  |  | Competitive condition<br>(N = 66) |  |  | Non-competitive condition |  |
| --- | --- | --- | --- | --- | --- | --- |
|  |  |  |  |  | Non-dyed food<br>(N = 19) | Dyed food<br>(N = 19) |
| Resident began to reproduce |  | 49 |  |  | 10 | 10 |
| Outcome | Category <sup>a)</sup><br>Winner | A<br>resident | B<br>Intruder | C<br>undecidable | – | – |
|  |  | 40 | 8 | 1 |  |  |
| Color of eggs | Only normal | 19 | 2 | – | 10 | 0 |
|  | Only pink | 1 | 1 | – | 0 | 10 |
|  | Normal and pink | 13 | 3 | – | 0 | 0 |
|  | No eggs | 7 | 2 | – | 0 | 0 |

**Table S3.**
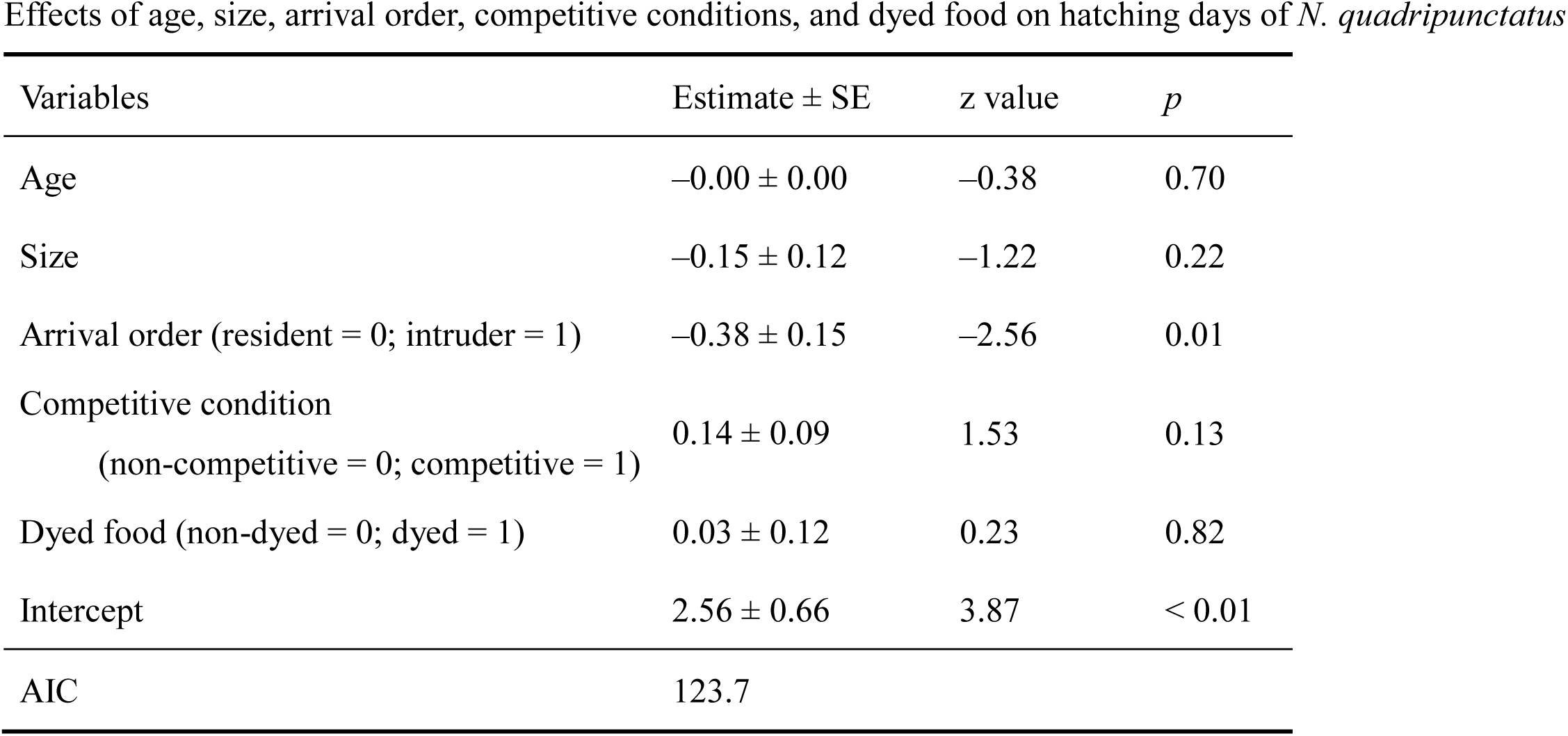
Effects of age, size, arrival order, competitive conditions, and dyed food on hatching days of *N. quadripunctatus*.

**Table S4.**
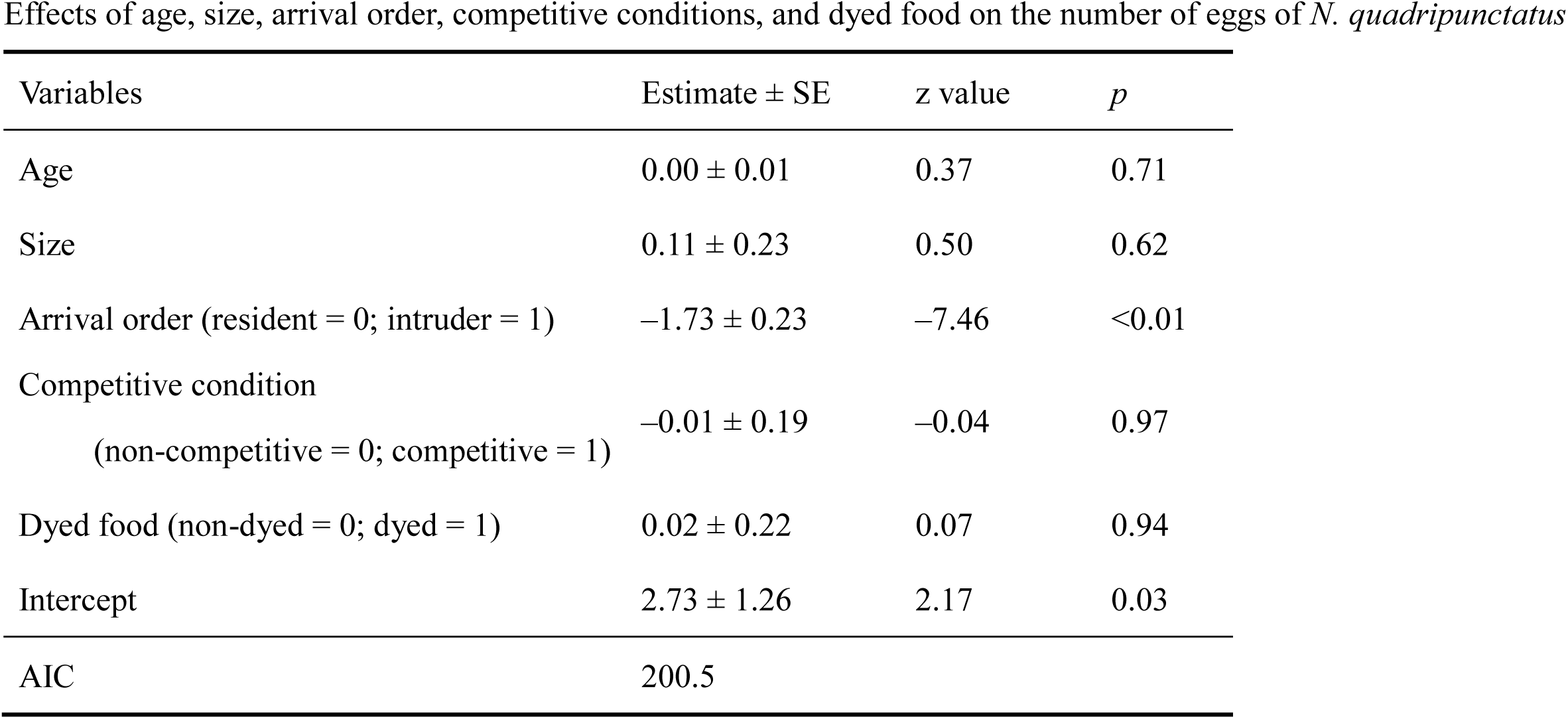
Effects of age, size, arrival order, competitive conditions, and dyed food on the number of eggs of *N. quadripunctatus*.

**Table S5.**
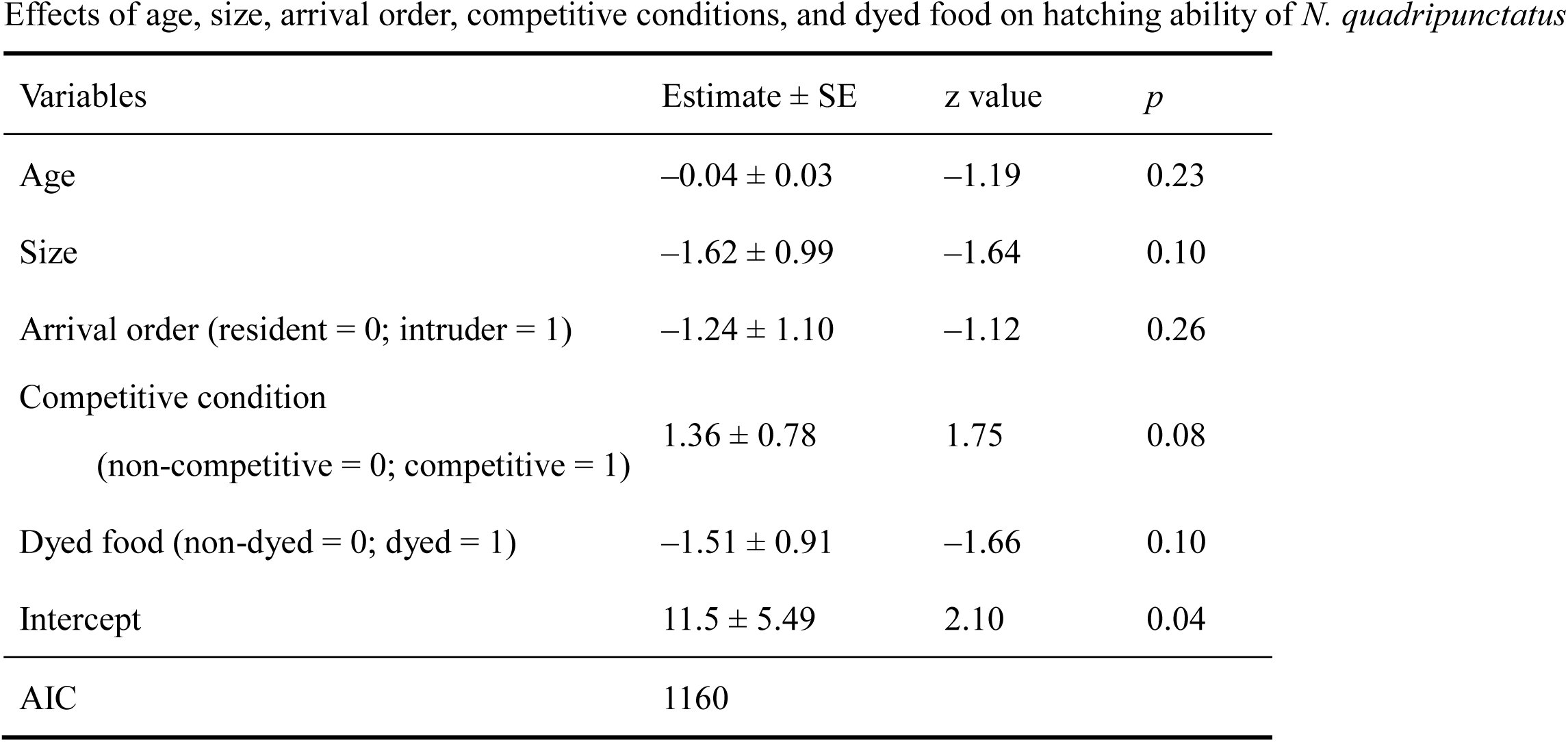
Effects of age, size, arrival order, competitive conditions, and dyed food on hatching ability of *N. quadripunctatus*.

| Variables | Estimate ± SE | z value | *p* |
| --- | --- | --- | --- |
| Age | $-0.04 \pm 0.03$ | -1.19 | 0.23 |
| Size | $-1.62 \pm 0.99$ | -1.64 | 0.10 |
| Arrival order (resident = 0; intruder = 1) | $-1.24 \pm 1.10$ | -1.12 | 0.26 |
| Competitive condition<br>(non-competitive = 0; competitive = 1) | $1.36 \pm 0.78$ | 1.75 | 0.08 |
| Dyed food (non-dyed = 0; dyed = 1) | $-1.51 \pm 0.91$ | -1.66 | 0.10 |
| Intercept | $11.5 \pm 5.49$ | 2.10 | 0.04 |
| AIC | 1160 |  |  |

**Table S6.** Effects of age, size, arrival order, competitive conditions, and dyed food on the egg volume of *N. quadripunctatus*.

| Variables | Estimate ± SE | t value | *P* |
| --- | --- | --- | --- |
| Age | 0.00 ± 0.01 | 0.92 | 0.37 |
| Size | -0.24 ± 0.16 | -1.45 | 0.16 |
| Arrival order (resident = 0; intruder = 1) | -0.44 ± 0.19 | -2.26 | 0.03 |
| Competitive condition<br>(non-competitive = 0; competitive = 1) | 0.56 ± 0.14 | 4.03 | <0.01 |
| Dyed food (non-dyed = 0; dyed = 1) | -0.07 ± 0.18 | -0.37 | 0.71 |
| Intercept | 2.72 ± 0.91 | 3.00 | 0.01 |
| AIC | Not available |  |  |

**Table S7.** Effects of resident age, intruder age, resident size, intruder size, and intruder oviposition on competitive duration of *N. quadripunctatus*.

| Variables | Estimate ± SE | z value | *p* |
| --- | --- | --- | --- |
| Resident age | $-0.00 \pm 0.02$ | -0.10 | 0.92 |
| Intruder age | $0.01 \pm 0.02$ | 0.35 | 0.72 |
| Resident size | $0.10 \pm 0.83$ | 0.12 | 0.91 |
| Intruder size | $-0.33 \pm 0.42$ | -0.79 | 0.43 |
| Intruder oviposition (no = 0; yes = 1) | $0.35 \pm 0.25$ | 1.42 | 0.16 |
| Intercept | $1.54 \pm 4.70$ | 0.33 | 0.74 |
| AIC | 29.27 |  |  |

**Table S8.** Effects of resident age, intruder age, resident size, and intruder size on intruder oviposition of *N. quadripunctatus*.

| variables | Estimate ± SE | z value | *p* |
| --- | --- | --- | --- |
| Resident age | $-0.04 \pm 0.06$ | -0.62 | 0.54 |
| Intruder age | $0.06 \pm 0.06$ | 0.96 | 0.34 |
| Resident size | $-1.78 \pm 2.65$ | -0.67 | 0.50 |
| Intruder size | $-2.49 \pm 1.47$ | -1.69 | 0.09 |
| Intercept | $21.9 \pm 15.2$ | 1.44 | 0.15 |
| AIC | 57.78 |  |  |

**Table S9.** Effects of age, size, arrival order, competitive conditions, and dyed food on the modified hatching day of *N. quadripunctatus*.

| variables | Estimate ± SE | z value | *p* |
| --- | --- | --- | --- |
| Age | $-0.00 \pm 0.00$ | -0.39 | 0.70 |
| Size | $-0.14 \pm 0.12$ | -1.22 | 0.22 |
| Arrival order | $-0.02 \pm 0.14$ | -0.15 | 0.88 |
| Competitive condition | $0.14 \pm 0.09$ | 1.57 | 0.12 |
| Dyed food | $0.03 \pm 0.12$ | 0.26 | 0.79 |
| Intercept | $2.52 \pm 0.64$ | 3.94 | < 0.01 |
| AIC | 124.2 |  |  |

In this analysis, two days were added to the hatching day of the intruders, which is the period from the first day of reproduction by residents to the hatching day of the intruders.

